# Investigation of the quorum-sensing regulon of the biocontrol bacterium *Pseudomonas chlororaphis* strain PA23

**DOI:** 10.1101/854596

**Authors:** Nidhi Shah, April Gislason, Michael Becker, Mark F. Belmonte, W.G. Dilantha Fernando, Teresa R. de Kievit

**Affiliations:** Department of Microbiology, University of Manitoba, Winnipeg, Manitoba, Canada; Department of Biological Sciences, University of Manitoba, Winnipeg, Manitoba, Canada; Department of Plant Science, University of Manitoba, Winnipeg, Manitoba, Canada

## Abstract

*Pseudomonas chlororaphis* strain PA23 is a biocontrol agent capable of protecting canola from stem rot disease caused by the fungal pathogen *Sclerotinia sclerotiorum*. PA23 produces several of inhibitory compounds that are under control of a complex regulatory network. Included in this cascade is the PhzRI quorum sensing (QS) system, which plays an essential role in PA23 biocontrol. The focus of the current study was to employ RNA sequencing to explore the spectrum of PA23 genes under QS control. Transcriptomic profiling revealed 545 differentially expressed genes (365 downregulated; 180 upregulated) in the *phzR* mutant and 534 genes (382 downregulated; 152 upregulated) in the AHL-deficient PA23-6863. In both strains, decreased expression of phenazine, pyrrolnitrin, and exoprotease biosynthetic genes was observed. We have previously reported that QS activates expression of these genes and their encoded products. In addition, elevated siderophore and decreased chitinase gene expression was observed in the QS-deficient stains, which was confirmed by phenotypic analysis. Inspection of the promoter regions revealed the presence of “*phz*-box” sequences in only 58 of the 807 differentially expressed genes, suggesting that much of the QS regulon is indirectly regulated. Consistent with this notion, 41 transcriptional regulators displayed altered expression in one or both of the QS-deficient strains. Collectively, our findings indicate that QS governs expression of approximately 13% of the PA23 genome affecting diverse functions ranging from secondary metabolite production to general metabolism. To the best of our knowledge, this represents the first global transcriptomic analysis of the QS regulon of a biocontrol pseudomonad.

## Introduction

Certain pseudomonads are able to inhibit fungal pathogens via the production of secondary metabolites through a process known as biological control. *Pseudomonas chlororaphis* strain PA23 is one such organism that suppresses canola stem rot caused by the fungal pathogen *Sclerotinia sclerotiorum* (1, 2). We have established that biocontrol by this bacterium occurs through direct and indirect mechanisms. Direct pathogen inhibition results from exposure to secreted bacterial products including the antibiotics pyrrolnitrin (PRN) and phenazine (PHZ), together with HCN, chitinases, proteases, lipases and siderophores (1, 3). PA23 also exerts its effects indirectly through priming the plant defense response, enabling the plant to more effectively fend off pathogen attack (4).

A complex regulatory network governs expression of PA23 antifungal (AF) compounds. At the top of this hierarchy sits the GacS-GacA two component signal transduction system that is essential for PA23 biocontrol (5, 6). Working in concert with Gac is the Rsm system, which consists of RsmA-like translational repressor proteins and small regulatory RNAs (7). Additional regulators overseeing production of PA23 biocontrol metabolites include the stationary phase sigma factor RpoS (8), PsrA (<u>P</u>seudomonas <u>S</u>igma <u>R</u>egulator <u>A</u>) (6), the stringent response (SR) (8), the anaerobic regulator ANR (9), and a novel LysR-type regulator called PtrA (10).

Adding to this complexity, PA23 AF compounds are expressed in a population-density-dependent fashion through quorum sensing (QS) (11). Like other gram-negative bacteria, PA23 uses N-acylhomoserine lactones (AHLs) as indices of population density (11–13). The first QS system identified in PA23 consists of the transcriptional regulator PhzR and the AHL synthase PhzI. The genes encoding these elements, *phzR* and *phzI*, are situated upstream of the *phzABCDEFG* biosynthetic locus responsible for PHZ production (11). Characterization of a *phzR* mutant (PA23*phzR*) and an AHL-deficient strain (PA23-6863) revealed a lack of fungal inhibition, which was attributed to reduced PHZ, PRN and protease production (11). A second QS system called CsaRI (<u>C</u>ell <u>S</u>urface <u>A</u>lteration) has been identified in the closely related *P. chlororaphis* 30-84 (14). The Csa system is not involved in the regulation of secondary metabolites or biocontrol genes, instead, it controls cell surface properties and biofilm formation (14). While homologs of *csaI* and *csaR* are present in the PA23 genome, the role of this QS system in PA23 remains unknown. Another QS system, called AurRI has been reported in *P. chlororaphis* subsp. *aurantiaca* PB-St2; however, it has not been characterized (15).

In *Pseudomonas aeruginosa*, global transcriptomic analysis using microarrays revealed that over 300 genes are under QS control (16, 17), greatly exceeding the number identified through more targeted approaches. For the first time, the magnitude of QS control was recognized to extend well beyond virulence, regulating diverse aspects of *P. aeruginosa* physiology. In organisms that employ AHL-based QS, control is mediated in one of two ways: directly through interaction with the promoter regions of target genes, or indirectly through other regulators. For several genes in the former category, consensus sequences have been identified that are required for LuxR-AHL complex binding (18). These “lux box-like” sequences are located in different positions depending on the gene in question (17, 18). In strain PA23, a “*phz*-box” sequence was identified upstream of the PHZ-biosynthetic locus as well as other genes under QS control (6, 11). We have previously established that the Phz QS system is deeply enmeshed in the complex hierarchy of gene regulation in strain PA23. Analysis of *gac* mutants revealed a lack of AHL production; consequently QS appears to be under control of this global regulatory system (5). The PhzRI QS system is also interconnected with RpoS (11), ANR (9), and the transcriptional regulator PtrA (19). As such, a large number of QS-regulated genes are expected to be indirectly regulated in this bacterium.

The focus of the current study was to explore the scope of genes under QS control in strain PA23 through RNA sequencing (RNA-seq). Analysis of a *phzR* mutant and an AHL-deficient derivative revealed that QS regulates approximately 13% of the PA23 genome, impacting diverse aspects of physiology ranging from secreted exoproducts to central metabolism. Moreover the vast majority of genes in the QS regulon appear to be indirectly controlled as very few contained *phz* boxes in their promoter regions. To the best of our knowledge, this is the first study to define the spectrum of genes under QS regulation in a biocontrol pseudomonad.

## Materials and Methods

### Bacterial strains and growth conditions

Bacterial strains used in this study are outlined in Table 1. *E. coli* strains were cultured on Lennox Luria Bertani (LB) media (Difco Laboratories, Detroit, MI) at 37°C. *P. chlororaphis* PA23 was cultured and maintained on LB media (Difco) at 28°C. *S. sclerotiorum* was cultured and maintained on Potato Dextrose Agar (PDA; Difco) at 22°C. Antibiotics were purchased from Research Products International Corp. (Prospect, IL) and supplemented at the following concentrations: gentamicin (Gm; 20 µg mL^−1^), tetracycline (Tc; 15 µg mL^−1^) for *P. chlororaphis* PA23 and Gm (15 µg mL^−1^) and Tc (15 µg mL^−1^) for *E. coli*. For phenotypic assays and cDNA library synthesis, strains were cultured in M9 Minimal Salts Media (M9; Difco) supplemented with 1 mM MgSO_4_ and 0.2% glucose, from here on referred to as M9-Glc.

**Table 1.**
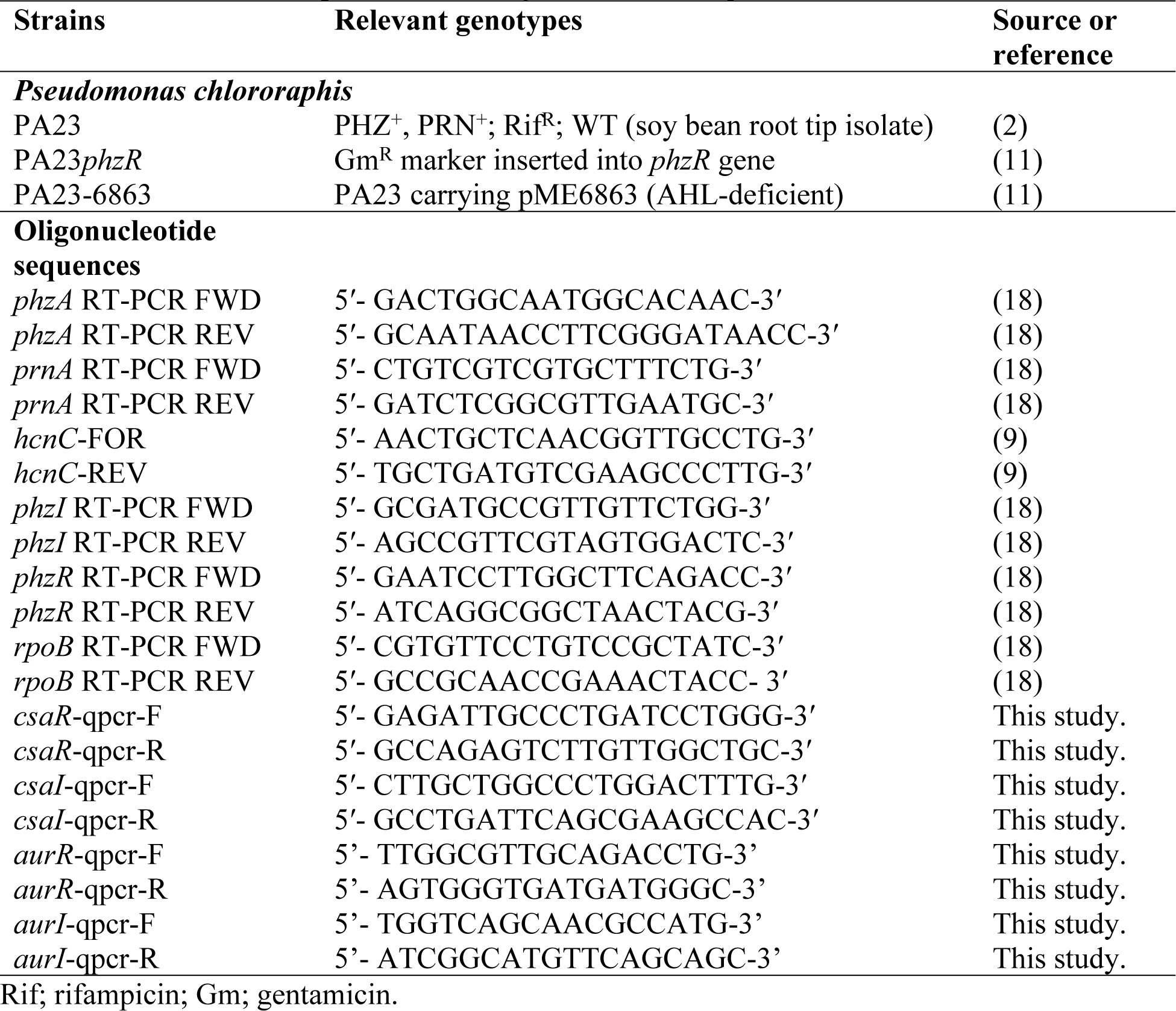
Bacterial strains, plasmids and oligonucleotides sequences

### Nucleic acid manipulations

Isolation, purification, endonuclease digestion and all other manipulation of DNA was performed according to the protocols described in Sambrook *et al.* (20). Polymerase Chain Reaction (PCR) was conducted following standard conditions outlined by New England Biolabs (NEB) (Ipswich, USA) data sheets supplied with their buffer system and *Taq* polymerase.

### RNA extraction and cDNA library synthesis

cDNA libraries were generated for PA23, PA23*phzR*, and PA23-6863. Three biological replicates of each strain were cultured in 30 ml of M9-Glc. Cells were harvested at early stationary phase (OD_600_1.20-1.50) by pelleting for 10 min at 6000 rpm at 4°C followed by flash freezing in liquid nitrogen. Pellets were stored at −80°C for up to one week. Total RNA was extracted using the Fermentas Plant RNA extraction kit (Waltham, USA) per manufacturer’s instructions. Residual genomic DNA was removed by treatment with TURBO RNAase-free DNAse I (Ambion, Carlsbad, USA) according to manufacturer’s instructions. RNA concentration was verified using a NanoVue spectrophotometer (GE Healthcare), and quality was measured using an Agilent 2100 Bioanalyzer with Agilent RNA 6000 Pico and Nano Chips (Agilent Technologies; Santa Clara, CA, USA). cDNA libraries were constructed using the alternative HTR protocol described by Kumar et al. (21) adapted for bacterial RNA. Ribosomal RNA was depleted using the MicrobExpress kit (Ambion) as per manufacturer’s instructions. Fragmentation time was reduced to 10 min and the number of cycles for final PCR amplification of the libraries was adjusted to 12 (21). Final cDNA libraries with ligated adaptors were size-selected to fall between 250 and 500 bp using the E-Gel® electrophoresis system (Invitrogen). cDNA quantity was measured using the Quant-iT™ PicoGreen® dsDNA Assay Kit (Thermo Fisher, Rockford, USA) with a Nanodrop 3300 (Thermo Fisher). cDNA was validated using Agilent Bioanalyser High Sensitivity DNA Chips (Agilent Technologies) at three points: i) after first and second strand cDNA synthesis; ii) after the final PCR amplification of the libraries; and iii) after size selection with the E-Gel® electrophoresis system. 100-bp single-end RNA-sequencing was carried out at Génome Québec (Montreal, Canada) on the Illumina HiSeq 2000 platform with a multiplex value of 9.

### Data analysis

Sequenced reads were analysed by FastQC to determine quality (http://www.bioinformatics.babraham.ac.uk/projects/fastqc/) and the Trimmomatic tool (22) enabled removal of low quality reads (Q < 30) and barcode adapters. Reads mapping to each gene was counted after mapping raw reads to the genome via the Rockhopper program (23) using the *P. chlororaphis* PA23 reference annotation from NCBI (gi: accession no. NZ_CP008696). Of total reads, 93-96% mapped to the *P. chlororaphis* PA23 genome across samples (S1Table). The R Bioconductor package DESeq2 (24) was employed to normalise raw read counts from each replicate and to determine significantly differentially expressed genes (adjusted p-value ≤ 0.01). This output enabled generation of gene expression profiles for each strain. The log_2_ fold change values ≥1.5 or ≤ −1.5 were selected based on previous analysis carried out by Shemesh *et al.*, (25) on the *Streptococcus mutans* transcriptome. Functional analysis was carried out via <u>C</u>luster of <u>O</u>rthologous <u>G</u>roups (COG) analysis. COG categories were assigned to the translated transcript sequences through the <u>C</u>onserved <u>D</u>omain <u>D</u>atabase (CDD) and the batch web-CD search tool (26, 27).

### Autoinducer assay

The amount of AHL produced by each strain was approximated by spotting 5 μl of an M9-Glc culture, grown in for 24 h and adjusted to an OD_600nm_ of 1, onto *Chromobacterium violaceum* CVO26 seeded LB agar plates. The radius of the purple zone surrounding each colony was measured after 24 h of incubation at 28°C.

### Siderophore assay

To analyze siderophore production, a 5µl volume of overnight bacterial culture grown in M9-Glc was spotted onto Chrome Azurol S (CAS) media (28). The diameter of the yellow zone surrounding the colonies, indicative of siderophore production, was measured following 24 hours of incubation at 28°C.

### Chitinase assay

Strains were tested for their ability to produce chitinase according to the protocol outlined by Wirth & Wolf (29). Strains were cultured in M9-Glc broth until they reached early stationary phase (OD_600_ 1.20-1.50). A 250-µl aliquot of cell free supernatant was incubated with equal parts of 0.1M NaOAc, pH 5.2 (250 µL) and carboxymethyl-chitin-Remazol brilliant violet aqueous solution (250 µL) (Blue Substrates, Göttingen, Germany). After incubation for 1 hour at either 28°C or 37°C, reactions were stopped by the addition of 250 µl of 1M HCl. Mixtures were cooled on ice for 10 min, spun at 20,000xg for 10 min and absorbance readings were taken at 550nm. Three replicates were analysed for each strain and experiments were performed in triplicate.

### Motility assays

Swimming motility assays were carried out by growing cultures of PA23 and derivative strains were to early stationary phase (24 h) in 3 ml of M9-Glc at 28°C. The cultures were then standardized to an OD_600nm_ of 1 and stab-inoculated (half-way down) into the centre of M9-Glc media solidified with 0.3% agar. Plates were incubated for 72 h at 28°C and the diameter of the swim zone was measured every 24 h. Five replicates were inoculated for each strain and three independent experiments were performed.

To assess swarming motility, fresh colonies of PA23 and derivative strains were gently inoculated using an applicator stick onto the surface of a swarm media plate (0.5% peptone, 0.3% yeast extract solidified with 0.8% agar). Plates were incubated for 96 h at 28°C, pictures of each plate were taken every 24 h and the area of the swarming colony was measured using ImageJ software (30). Five plates were analyzed for each strain and three independent experiments were performed.

### Quantitative PCR (qPCR)

Quantitative PCR (qPCR) was used to monitor expression of metabolite and regulatory genes involved in biocontrol. Triplicate cultures of PA23 and derivative strains were grown to early stationary phase in a 3ml volume of M9-Glc. Cells were harvested by incubating 500µl of culture with 2x volume of RNAprotect reagent (QIAGEN, Valencia, USA) for 5 min followed by centrifugation for 10 min at 6000 rpm. Pellets were stored at −20° for up to one week. Total RNA was extracted using the RNeasy Mini Kit (QIAGEN). Residual genomic DNA was removed by treatment with TURBO RNAase-free DNAse I (Ambion). RNA concentrations were measured at absorbance 260 and 280 nm; only RNA samples with A260/A280 between 1.8 and 2.0 were used in subsequent steps. cDNA was generated by reverse transcription using the Maxima First Strand cDNA Synthesis Kit (Thermo Scientific) and random hexamer primers in a 20-µL total reaction volume. The following conditions were employed: initial heating at 25°C for 10 min, reverse transcription at 50°C for 15 min, and enzyme denaturation at 85°C for 5 min. Sequences for the PA23 genes of interest were obtained from GenBank (gi: NZ_CP008696) and the primer sequences are listed in Table 1. PCR was performed using a CFX96 Connect^TM^ Real-Time PCR Detection System (Bio-rad, Hercules, USA) and SsoFast^TM^ EvaGreen® Supermix (Bio-rad). The final 10-µL volume mixture in each well contained 0.4 µL of both forward and reverse primers (12µM), 1 µL of 1:20 diluted cDNA, 5 µL of SsoFast^TM^ EvaGreen® Supermix and 3.4 µL of nuclease-free water. PCR reaction conditions included an initial denaturation at 98°C for 2 min, followed by 39 cycles of 98°C for 5 s, 60°C for 30 s, and 60◦C for 5 s. Melt-curve analysis was performed to evaluate the formation of primer dimers and other artefacts to validate results. Each reaction was performed in triplicate and experiments were repeated three times with three biological replicates. Relative gene expression was calculated using the ΔΔCt method as described by Livak & Schmittgen (31), with *rpoB* as the reference gene and the CFXManager^TM^ software (Bio-rad).

### *phz-*box identification

The presence of putative *phz*-boxes upstream of differentially expressed genes determined through RNAseq were identified using the MEME suite Motif Alignment and Search Tool (MAST) algorithm (http://meme-suite.org/tools/mast) (32, 33). A *phz-*box consensus sequence VCKRCHWGHKYKBSHWRK, (where V= A, C or G; K= G, T; R= A, G; H= C, T or A; W= A, T; Y=C, T; B= C, G or T; S= G, C; N= A, T, G or C) was generated using previously identified *phz*-boxes in PA23 (9, 11). Only consensus sequences located within 500 basepairs upstream of the translational start site were considered significant.

## Results and Discussion

### Identification of genes under QS control

In PA23, the Phz QS system is essential for biocontrol as strains lacking this regulatory circuitry no longer produce the antibiotics and degradative enzymes required for fungal antagonism (11). To fully appreciate the global effect of QS on PA23 gene expression, RNAseq was conducted on two QS-deficient strains. The *phzR* mutant, PA23*phzR*, was generated through allelic exchange and exhibits reduced AHL levels compared to wild type (S1 Fig.) (11). Conversely, PA23-6863 contains the *B. subtilis* AHL lactonase-encoding *aiiA* gene on a plasmid and is AHL-deficient (S1 Fig.) (11). Because the PA23 genome encodes three QS systems including PhzRI, CsaRI and AurRI, PA23-6863 is equivalent to a triple mutant. This is relevant because in the closely related *P. chlororaphis* subsp. *aurantiaca* strain PB-St2, the three AHL synthases, PhzI, CsaI and AurI, produce structurally similar molecules (15). The presence of the lactonase ensures that there is no cross-activation by non-cognate AHLs.

We chose M9-Glc media to simulate the nutrient-limiting conditions present in the environment and cells were harvested at early stationary phase (OD_600_ 1.2-1.5) when secondary metabolite production occurs. Five hundred and forty-five differentially expressed genes were identified in the *phzR* mutant background (365 downregulated; 180 upregulated), corresponding to 8.8% of the genome (S2 Table). In the AHL-deficient PA23-6863, a total of 534 genes showed altered expression (382 downregulated; 152 upregulated) representing 8.6% of the genome (S3 Table). In total, 807 of the 6,179 coding sequences in the PA23 genome showed differential expression in one or both of the QS-deficient strains.

The degree of overlap between differentially expressed genes in the *phzR* mutant and in the AHL-deficient strain is quite low; 36.8% of downregulated and 26.2% of upregulated genes were conserved between them (Fig. 1). Genes showing altered expression exclusively in PA23-6863 is likely due to the fact that in this strain, all signal molecules are degraded leading to a total loss of QS; conversely in PA23*phzR*, only the Phz system is affected. Intriguingly, 275 genes showed differential regulation in the *phzR* mutant but not the AHL-deficient strain. There are examples of LuxR proteins that regulate gene expression in the absence of AHL; for instance in *Pectobacterium atrosepticum*, VirR binds to the promoter of the *rsmA* gene activating transcription. Upon binding to 3-oxo-C6-HSL, VirR undergoes a conformation change that results in dissociation from the promoter and *rsmA* is no longer expressed (34–36). The *P. aeruginosa* transcriptional regulator RhlR was recently reported to regulate gene expression in a C4-HSL-dependent and C4-HSL-independent manner (37). In the absence of the AHL synthase (RhlI), RhlR directs expression of genes involved in biofilm formation and virulence factors (37). Finally, AHL-deficient strains of *Pseudomonas corrugata* and *Pseudomonas brassicacearum* are phenotypically similar to wild type; whereas loss of the LuxR homolog results in dramatically altered traits (38) (Saikai and de Kievit, unpublished data). Collectively, these findings support a regulatory role for LuxR proteins in the absence of AHL binding.

**Figure 1:**
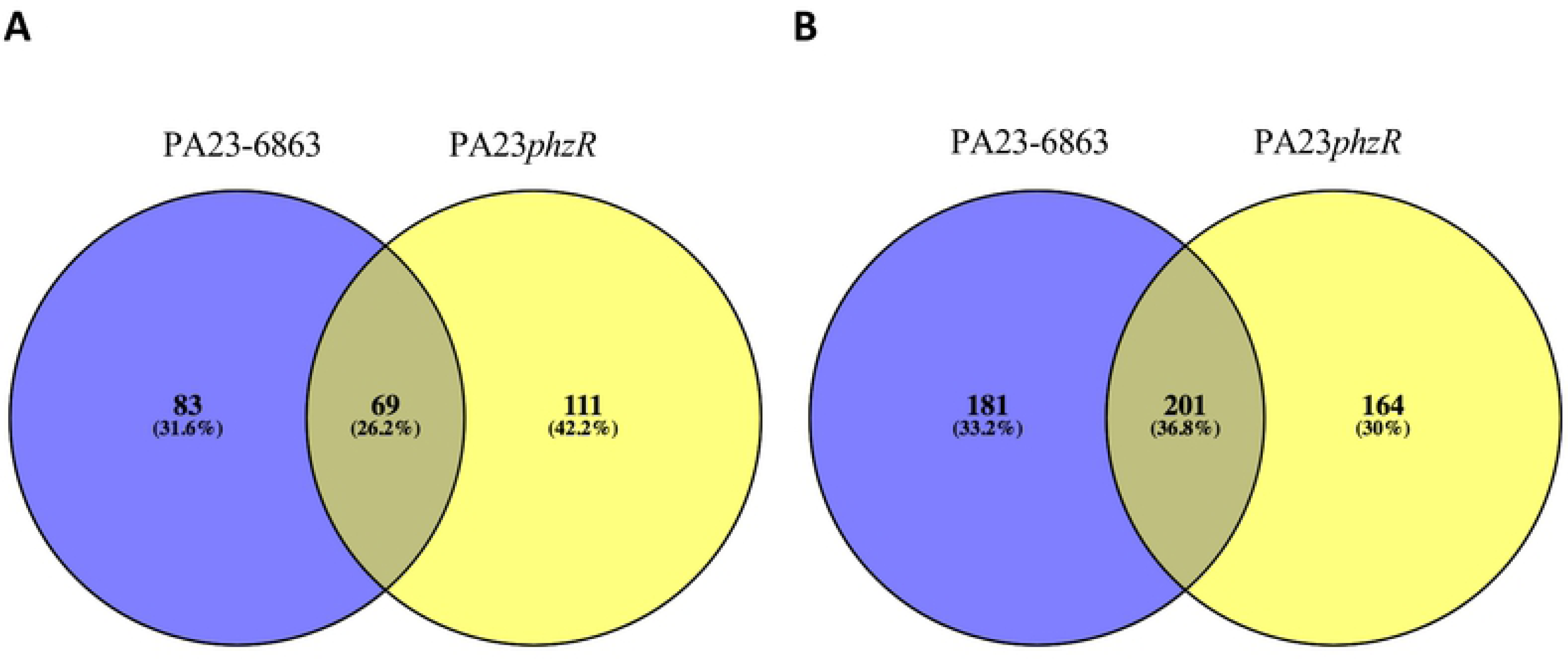
Comparison of the PA23*phzR* and PA23-6863 regulons. Venn diagrams were generated using Venny 2.1 (39) to determine the overlap of up (A) and down (B) regulated genes between PA23*phzR* and PA23-6863.

While the global impact of QS in a biocontrol pseudomonad has yet to be undertaken, microarray studies have been performed on two pathogenic pseudomonads, namely *P. aeruginosa* PAO1 and *P. syringae* B728a (16,17,40,41) Results from the current study show that expression of 13.06% of the PA23 genome is governed by QS. In *P. aeruginosa* PAO1, QS-modulated genes were reported to account for 6-10% of the genome and the majority of genes were positively regulated (16, 17). We discovered that in PA23*phzR*, 67% of the differentially regulated genes showed decreased expression, while 33% were upregulated. Similarly, in PA23-6863, 71.5% and 28.5% of genes displayed decreased and increased expression, respectively. Thus, for both PAO1 and PA23, QS is largely serving as a positive regulator of gene expression. In stark contrast to these large regulons, only 9 genes were found to be controlled by the *P. syringae* QS circuitry (41). All nine genes were located near the *ahlR* locus and all were positively regulated by QS.

### Functional characterisation of differentially expressed genes in PA23*phzR* and PA23-6863

Next, we sought to predict the functional role of the PA23 QS-controlled genes through <u>C</u>luster of <u>O</u>rthologous <u>G</u>roup (COG) analysis. COG clusters are constructed using functional characterization based on prokaryotic genomes (42). According to their predicted function, the 545 differentially expressed genes in the *phzR* mutant could be divided into 21 COG categories (Fig. 2a, S2a Fig., S2 Table). Similarly, the 534 genes identified in PA23-6863 could be grouped into 22 COG categories (Fig. 2b, S2b Fig., S3 Table). Several of the more relevant categories are discussed in detail in the following sections.

**Figure 2.**
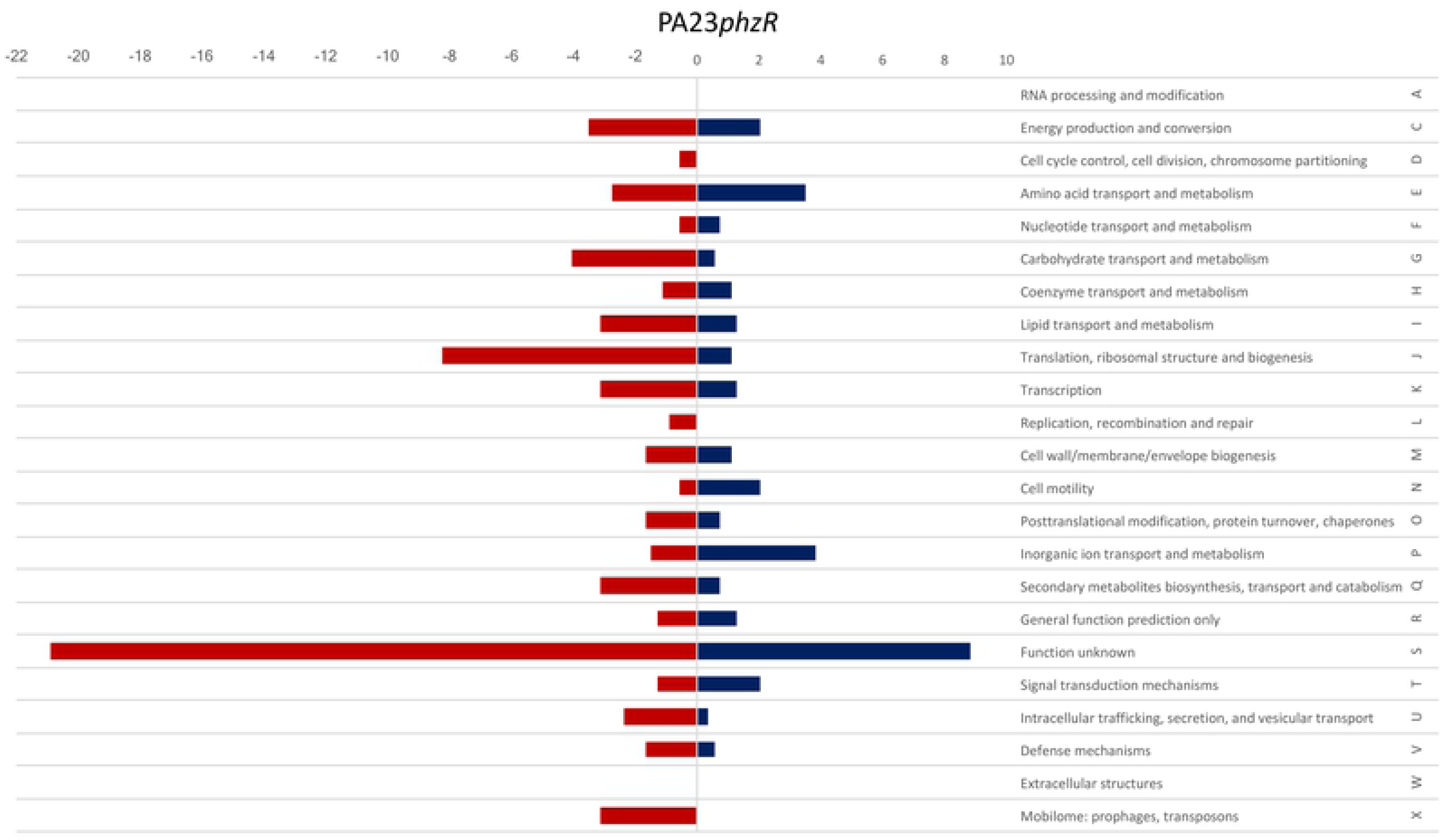
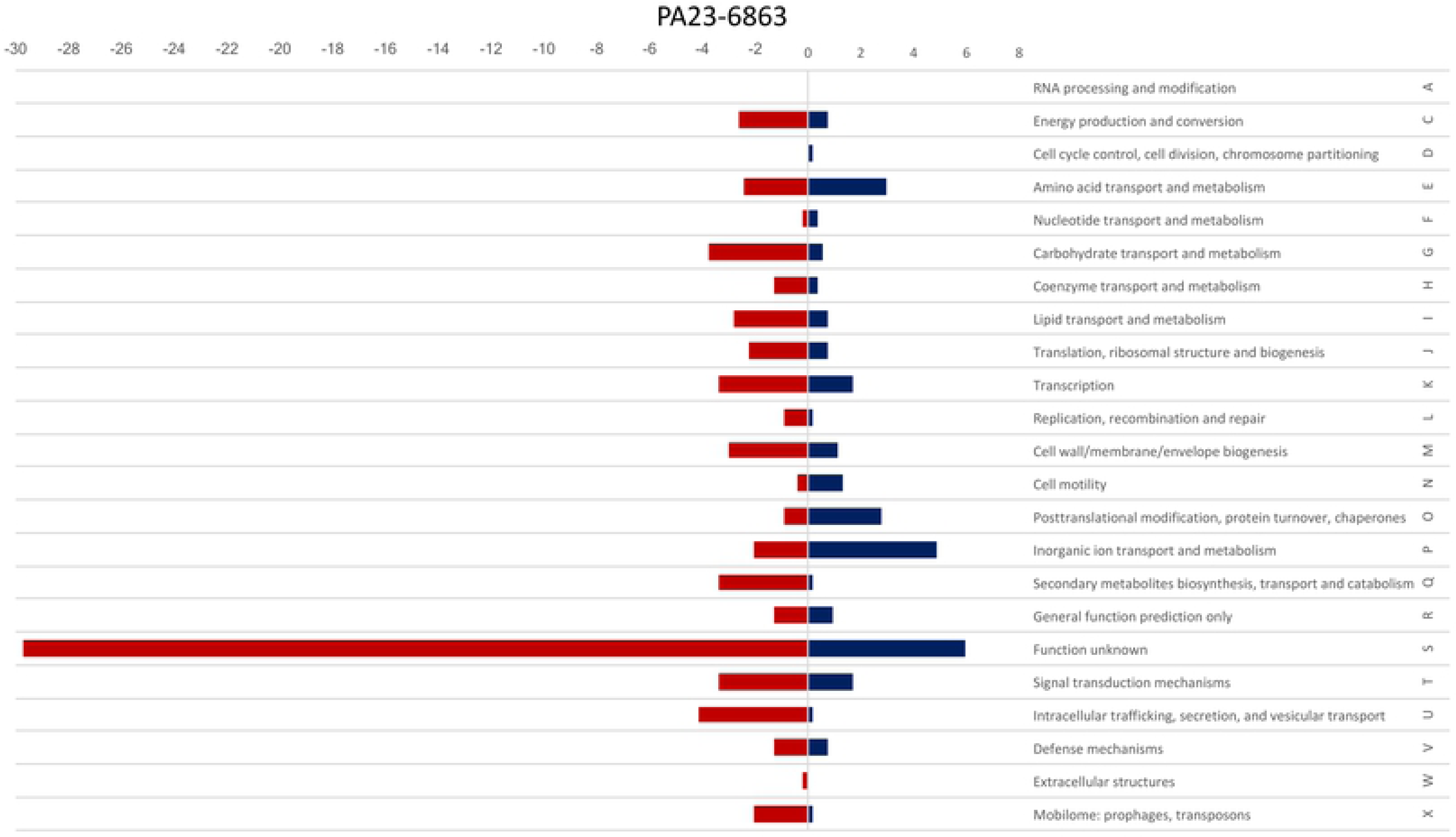
Functional analysis of differentially expressed genes in (A) PA23*phzR* and (B) PA23-6863 in comparison to wild type using Cluster of Orthologous Group (COG) analysis. The red bars indicate percent of differentially regulated genes that are downregulated, and the blue bars indicate percentage of genes that are upregulated in each category.

### Role of QS in regulation of PA23 secondary metabolites

In both QS-deficient strains, a number of genes involved in secondary metabolite production were significantly downregulated. For example, genes required for synthesis of PHZ (EY04_RS25715-45), hydrogen cyanide (EY04_RS11540-50), and exoprotease (EY04_RS11085) exhibited ≥3.42 log_2_fold decreased expression compared to the WT. Similarly, expression of the *prn* biosynthetic genes (EY04_RS17650-35) was reduced 1.93- and 4.17-log_2_fold in PA23*phzR* and PA23-6863, respectively. In a previous study, PA23 QS-deficient strains exhibited decreased *phzA-lacZ* and *prnA-lacZ* transcription and reduced PHZ, PRN and protease production (11), consistent with our RNAseq data. When qPCR was used to validate our global transcriptomic findings, expression of the *phz, prn* and *hcn* genes in PA23*phzR* and PA23-6863 was decreased (Fig. 3).

**Figure 3.**
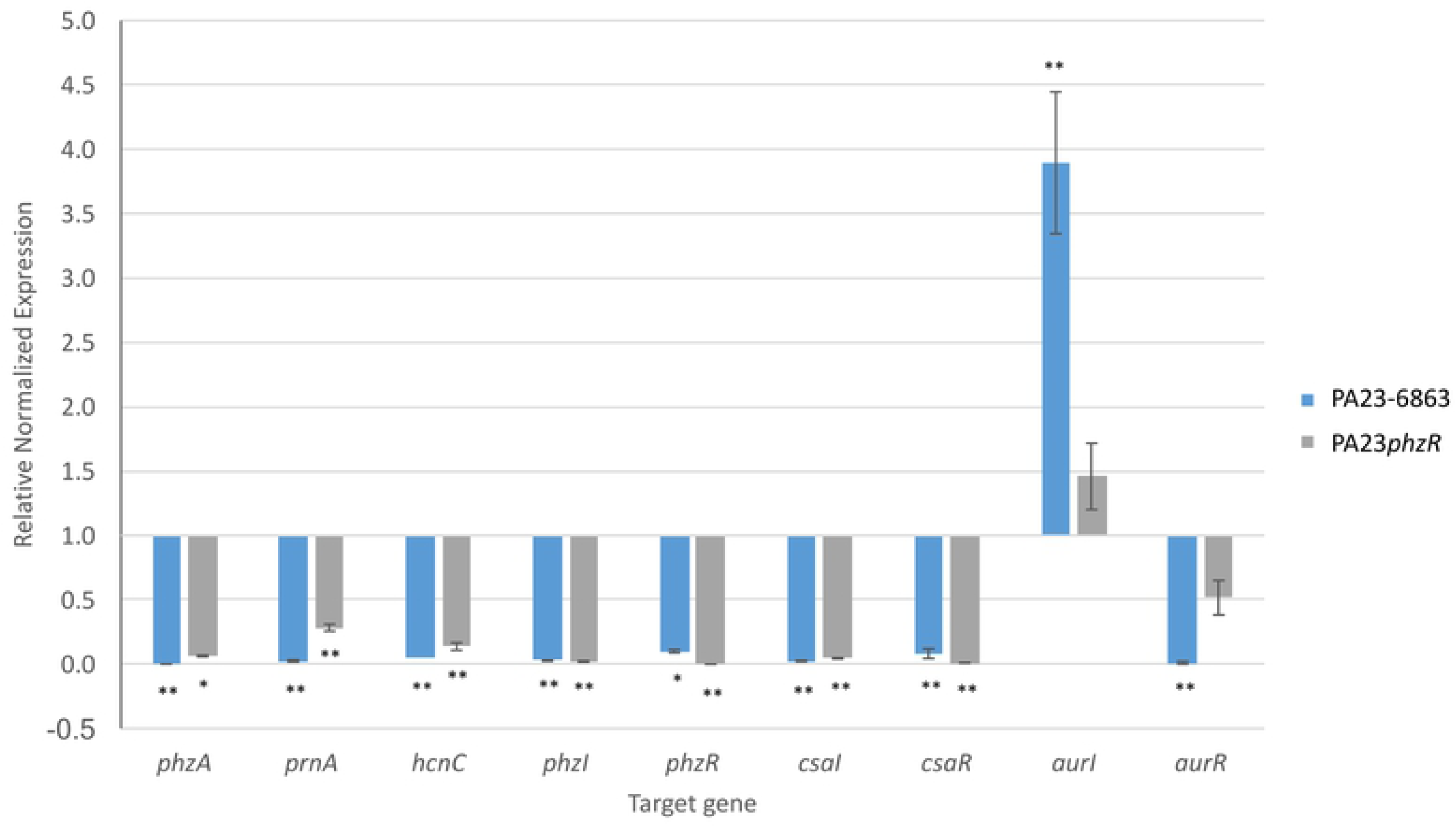
qPCR fold change in gene expression in PA23*phzR* and PA23-6863 compared to PA23 wild type. Analyzed genes were compared against *rpoB* as a reference gene. Gene expression in the wild type was normalized to 1.0. For strains that differ significantly from the wild type, columns have been marked with an asterisk (* < 0.01; **p < 0.001).

Other exoproduct genes that were differentially expressed in both QS derivatives include chitinase-encoding genes (EY04_RS16020, EY04_RS09705; downregulated ≥6.91-log_2_fold) and siderophore biosynthetic genes (EY04_RS15355-410; upregulated ≥ 2.61-log_2_fold). When we employed end-product analysis to support these findings, chitinase activity was completely abolished in both PA23*phzR* and PA23-6863 (0.00 ± 0.001) compared to the wild type (0.20 ± 0.006). On CAS agar, orange halos indicative of siderophore production, were significantly larger around colonies of PA23*phzR* (8.0±0.3 mm) and PA23-6863 (7.5±0.1 mm) compared to the PA23 parent (3.5±0.2 mm). Collectively, the RNAseq findings are in keeping with phenotypic characteristics of the QS-deficient strains, validating the robustness of this approach for defining the PA23 QS regulon (3, 11).

### Role of QS in motility

Several motility genes were significantly upregulated in the QS-deficient strains (S3 Fig., S2 Table and S3 Table). In addition to a number of chemotaxis signal transduction genes being induced, genes predicted to modulate cellular levels of cyclic di-GMP (EY04_RS05905, EY04_RS09010, EY04_RS10080, EY04_RS15460, EY04_RS24510) were upregulated in one or both of the QS-deficient strains. Notably, flagellin (EY04_RS07580), a diguanylate phosphodiesterase (EY04_RS05905), as well as the alternative stators, MotD (EY04_RS07755) and MotY (EY04_RS24205) were uniquely upregulated in PA23*phzR* (S3 Fig.). The MotCDY stators along with low levels of c-di-GMP are required to provide higher torque for flagellar movement through high agar concentrations in *P. aeruginosa* (43, 44). Such findings suggest that PA23*phzR* and PA23-6863 are more motile. Consistent with this, the swim zone of PA23*phzR* was significantly larger than the WT and PA23-6863, while PA23-6863 was more motile than the WT strain in 0.3% agar (Table 2, S3 Fig.). Since the swarming pattern of PA23 is irregular (5), the area of the swarming colony on 0.8% agar was measured using ImageJ software (30). After four days, PA23*phzR* swarmed significantly more than the WT strain (Table 2, S3 Fig.). PA23-6863 showed more variation between replicates and the area of swarming was not significantly different from the WT, but was less than PA23*phzR* (Table 2, S3 Fig.). These results are consistent with findings that QS suppresses motility in other rhizobacteria, such as *P. syringae* and *Sinorhizobium meliloti* (45–47). Since the switch from a sessile to a motile cell is energetically costly and flagella can induce plant defenses, it is beneficial for bacteria to tightly regulate motility until a lack of nutrients necessitate movement to a more nutritive environment (48).

**Table 2.** Motility of PA23 and derivative strains.

| Strain | Motility |  |  |  |
| --- | --- | --- | --- | --- |
|  | Swim 0.3% agar* |  |  | Swarm 0.8% agar |
|  | 24 h | 48 h | 72 h | 96 h† |
| PA23 WT | 7.2 (0.8) | 10.4 (1.1) | 12.4 (1.3) | 162.8 (18.6) |
| PA23*phzR* | 23.0 (2.1)§ | 46.6 (2.6)§ | 69.8 (3.6)§ | 815.2 (304.4)\|\| |
| PA23 -6863 | 11.3 (1.5)\|\| | 24.0 (1.0)§ | 35.1 (2.9)§ | 83.2 (50.1)# |

\*Mean (SD) diameter (mm) of the swim zone obtained from three independent experiments.
†Mean (SD) area (mm<sup>2</sup>) of swarming colony obtained from three independent experiments.
§Significantly different from WT (P<0.001).
||Significantly different from WT (P<0.05).

### Characterization of QS-regulated genes involved in other physiological processes

The largest percentage of QS regulated transcripts encoded proteins of unknown function (35.8% AHL^−^; 29.7% *phzR*^−^), which was also observed by Wagner et al. (17). Other major categories found to be QS regulated include those associated with cellular energetics and metabolism, such as energy production and conversion (category C; 3.4% AHL^−^; 5.5% *phzR*^−^), amino acid transport and metabolism (category E; 5.4% AHL^−^; 6.2% *phzR*^−^), carbohydrate transport and metabolism (category G; 5.4% AHL^−^; 4.6% *phzR*^−^), and lipid transport and metabolism (category I; 3.6% AHL^−^; 4.4% *phzR*^−^). Consistent with these findings, PAO1 genes linked to amino acid biosynthesis and metabolism, carbon compound catabolism, energy metabolism, and fatty acid and phospholipid metabolism were differentially expressed in the absence of QS (17). Genes involved in translation, ribosomal structure and biogenesis were also found to be under QS control (category J; 3.0% AHL^−^; 9.4% *phzR*^−^), including a large number of 50S and 30S ribosomal proteins as well as the 16S rRNA processing protein, RimM. Similar results were demonstrated for the PfsI/R QS regulon of the rice pathogen *Pseudomonas fuscovaginae*, where the majority of genes positively regulated by PfsI/R are involved in translation, ribosomal structure and biogenesis, including 20 ribosomal proteins and RimM (49).

In *P. aeruginosa*, QS controls expression of not only virulence factors but the secretion systems required for export (17). Our transcriptomic analysis revealed decreased expression of genes encoding type IV (EY04_RS00545) and VI (EY04_RS29490, EY04_RS29495, EY04_RS29520) secretion systems in one or both of the PA23 QS-deficient strains. Moreover, expression of resistance-nodulation-cell division (RND) efflux transporters including MexE homologues (EY04_RS19235, EY04_RS17230, EY04_RS17225, EY04_RS00160) was reduced in the absence of QS. Similarly in strain PAO1, three RND efflux systems were found to be under QS control (17). At present, the mechanism by which PA23 biocontrol metabolites are transported outside of cells has not been elucidated. We hypothesize that at least some of these compounds are exported via these secretion systems and/or active efflux.

### QS directly and indirectly regulates gene expression in PA23

For genes under QS control, regulation may occur in one of two ways: directly through PhzR-AI binding to the promoter or indirectly through control of other regulators. For those in the former category, an activated PhzR dimer is believed to bind to a highly-conserved consensus sequence known as the *phz*-box, previously identified upstream of the PA23 *phzA* and *phzI* genes (11). In *P. aeruginosa*, 7% of QS-regulated genes reportedly contain *las-* or *rhl-*boxes in the promoter region (16, 17). To determine if a similar trend is observed in PA23, we analysed the genome for *phz-*boxes using the Motif Alignment and Search Tool (MAST) algorithm (32, 33). In PA23, 545 and 534 genes were differentially regulated in the PhzR- and AHL-deficient backgrounds, respectively. However, only 58 genes contained *phz*-boxes within the 500-bp region upstream of the translational start (Table 3). A large number of transcriptional regulators display altered expression in the QS-deficient strains (Table 4). In total, 41 genes encoding TS regulators showed differential expression in one or both of the QS-deficient strains. The expression of 11 genes encoding regulators was altered in both strains, while 18 and 12 genes exhibited altered expression in only PA23-6863 and PA23*phzR*, respectively (Table 4). These findings support the hypothesis that a substantial proportion of the QS regulon is subject to indirect control. While a detailed discussion of all 41 regulators is not feasible, examples of prominent genes classified into each of these categories are provided below.

**Table 3.**
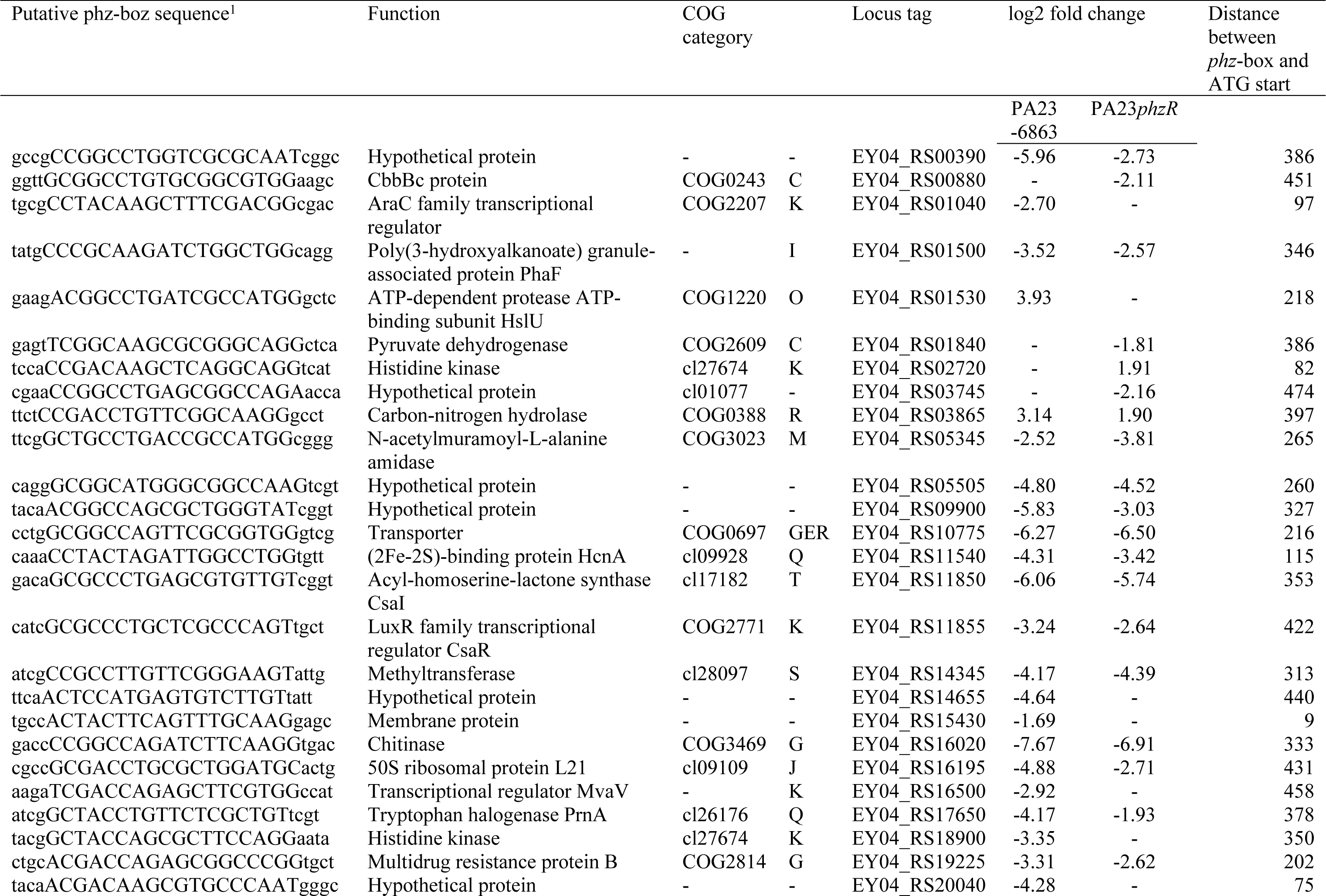
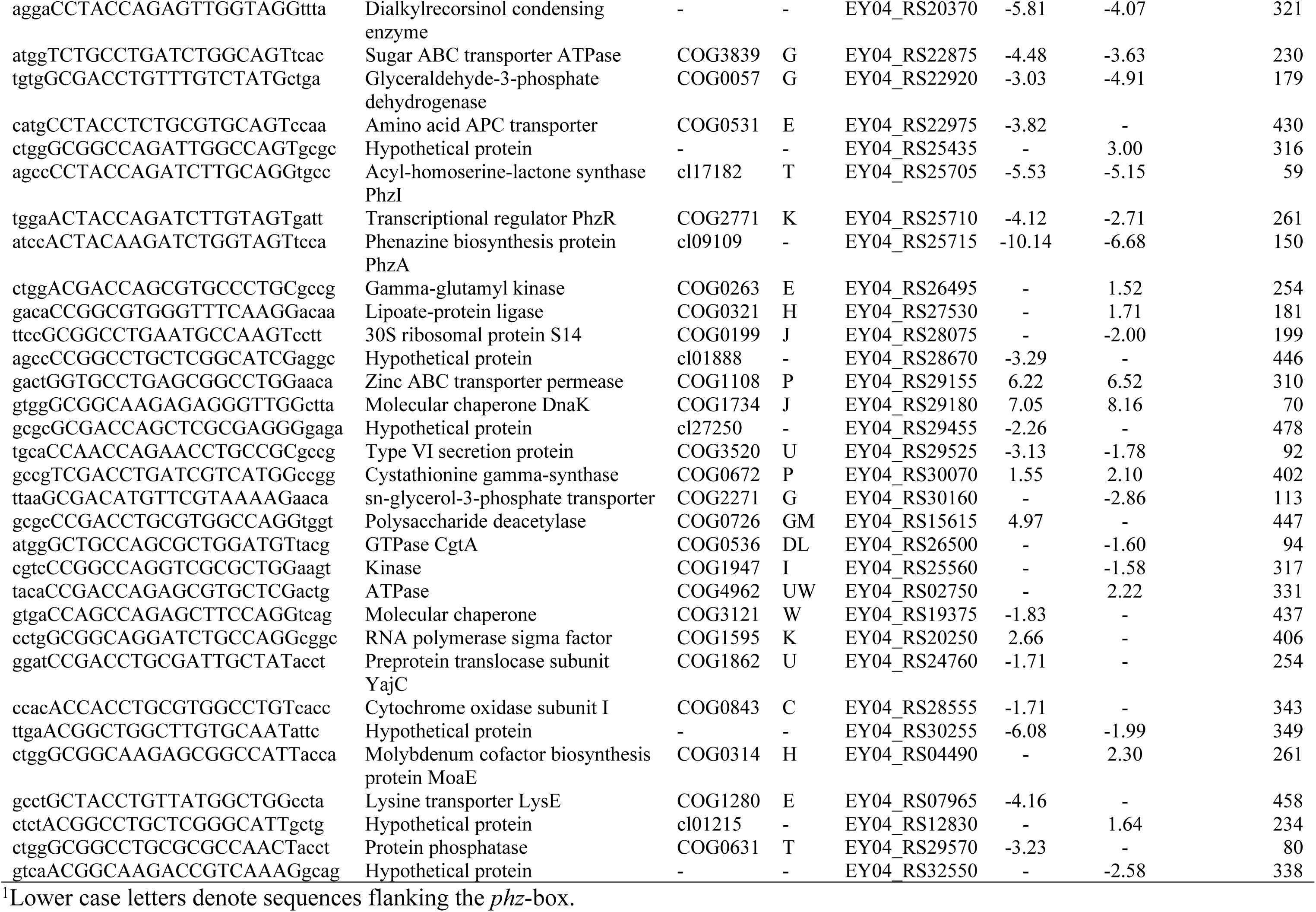
Genes containing a *phz*-box sequence within 500 bp of the ATG start

**Table 4.**
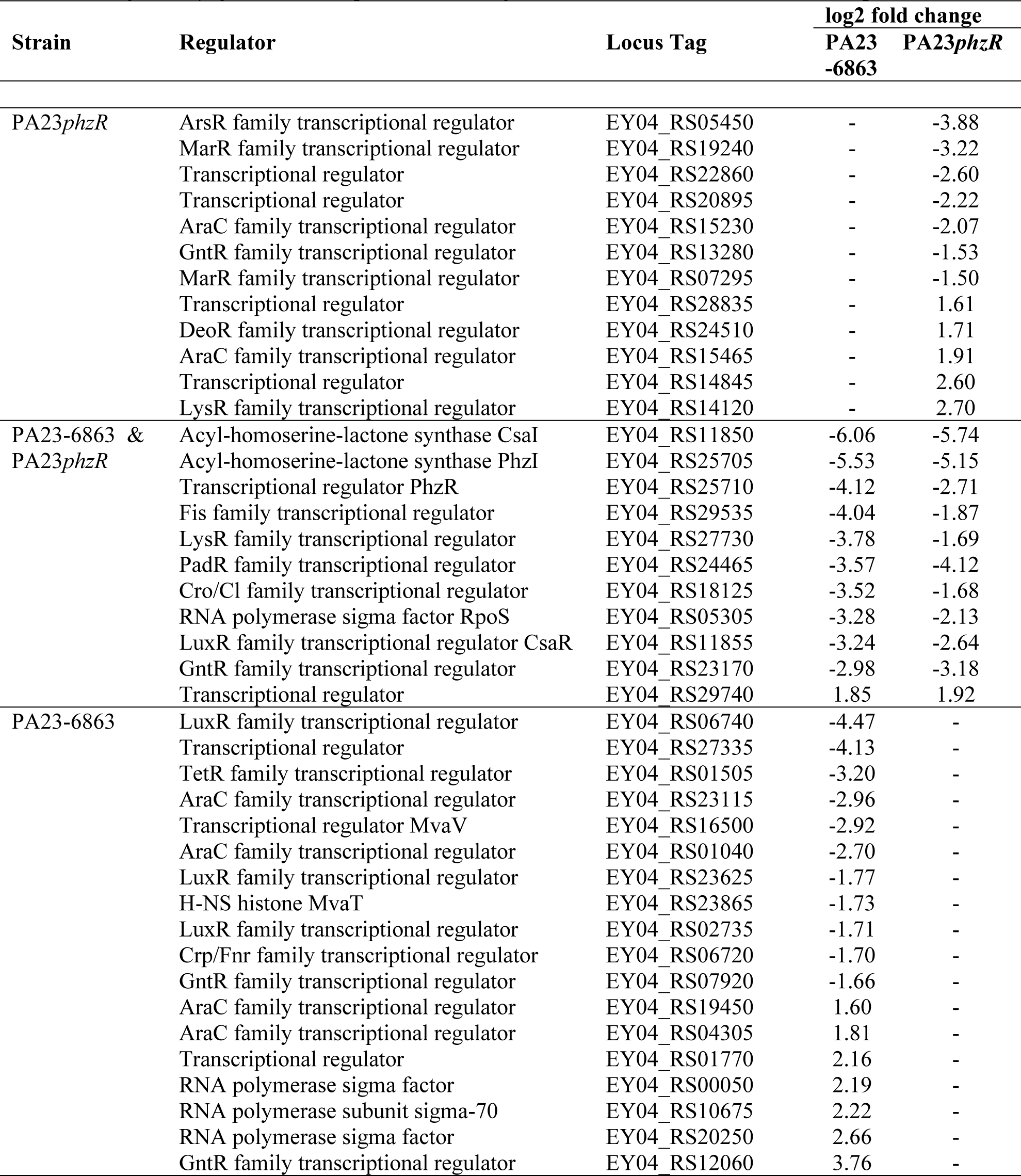
Regulatory genes under quorum-sensing control in *Pseudomonas chlororaphis* PA23.

| Strain | Regulator | Locus Tag | log2 fold change |  |
| --- | --- | --- | --- | --- |
|  |  |  | PA23<br>-6863 | PA23*phzR* |
| PA23*phzR* | ArsR family transcriptional regulator | EY04_RS05450 | - | -3.88 |
|  | MarR family transcriptional regulator | EY04_RS19240 | - | -3.22 |
|  | Transcriptional regulator | EY04_RS22860 | - | -2.60 |
|  | Transcriptional regulator | EY04_RS20895 | - | -2.22 |
|  | AraC family transcriptional regulator | EY04_RS15230 | - | -2.07 |
|  | GntR family transcriptional regulator | EY04_RS13280 | - | -1.53 |
|  | MarR family transcriptional regulator | EY04_RS07295 | - | -1.50 |
|  | Transcriptional regulator | EY04_RS28835 | - | 1.61 |
|  | DeoR family transcriptional regulator | EY04_RS24510 | - | 1.71 |
|  | AraC family transcriptional regulator | EY04_RS15465 | - | 1.91 |
|  | Transcriptional regulator | EY04_RS14845 | - | 2.60 |
|  | LysR family transcriptional regulator | EY04_RS14120 | - | 2.70 |
| PA23-6863 &<br>PA23*phzR* | Acyl-homoserine-lactone synthase CsaI | EY04_RS11850 | -6.06 | -5.74 |
|  | Acyl-homoserine-lactone synthase PhzI | EY04_RS25705 | -5.53 | -5.15 |
|  | Transcriptional regulator PhzR | EY04_RS25710 | -4.12 | -2.71 |
|  | Fis family transcriptional regulator | EY04_RS29535 | -4.04 | -1.87 |
|  | LysR family transcriptional regulator | EY04_RS27730 | -3.78 | -1.69 |
|  | PadR family transcriptional regulator | EY04_RS24465 | -3.57 | -4.12 |
|  | Cro/CI family transcriptional regulator | EY04_RS18125 | -3.52 | -1.68 |
|  | RNA polymerase sigma factor RpoS | EY04_RS05305 | -3.28 | -2.13 |
|  | LuxR family transcriptional regulator CsaR | EY04_RS11855 | -3.24 | -2.64 |
|  | GntR family transcriptional regulator | EY04_RS23170 | -2.98 | -3.18 |
|  | Transcriptional regulator | EY04_RS29740 | 1.85 | 1.92 |
| PA23-6863 | LuxR family transcriptional regulator | EY04_RS06740 | -4.47 | - |
|  | Transcriptional regulator | EY04_RS27335 | -4.13 | - |
|  | TetR family transcriptional regulator | EY04_RS01505 | -3.20 | - |
|  | AraC family transcriptional regulator | EY04_RS23115 | -2.96 | - |
|  | Transcriptional regulator MvaV | EY04_RS16500 | -2.92 | - |
|  | AraC family transcriptional regulator | EY04_RS01040 | -2.70 | - |
|  | LuxR family transcriptional regulator | EY04_RS23625 | -1.77 | - |
|  | H-NS histone MvaT | EY04_RS23865 | -1.73 | - |
|  | LuxR family transcriptional regulator | EY04_RS02735 | -1.71 | - |
|  | Crp/Fnr family transcriptional regulator | EY04_RS06720 | -1.70 | - |
|  | GntR family transcriptional regulator | EY04_RS07920 | -1.66 | - |
|  | AraC family transcriptional regulator | EY04_RS19450 | 1.60 | - |
|  | AraC family transcriptional regulator | EY04_RS04305 | 1.81 | - |
|  | Transcriptional regulator | EY04_RS01770 | 2.16 | - |
|  | RNA polymerase sigma factor | EY04_RS00050 | 2.19 | - |
|  | RNA polymerase subunit sigma-70 | EY04_RS10675 | 2.22 | - |
|  | RNA polymerase sigma factor | EY04_RS20250 | 2.66 | - |
|  | GntR family transcriptional regulator | EY04_RS12060 | 3.76 | - |

### Regulatory genes showing differential expression in both QS-deficient strains

*phzI* and *phzR* are among the 11 regulatory genes that showed differential expression in PA23*phzR* and PA23-6863 (Table 4). As expected, both genes were positively regulated consistent with the paradigm of autoinduction (Table 4). The gene encoding RpoS showed 2.13- and 3.28-log_2_fold lower expression in PA23*phzR* and PA23-6863 respectively. Cross-regulation between QS and RpoS has been previously demonstrated (11). *rpoS*, was found to be positively regulated by the Phz QS system (11), in keeping with findings presented herein.

Our RNAseq analysis revealed a connection between the Csa and Phz QS systems. In PA23*phzR* and PA23-6863, *csaR* was downregulated by a factor of 2.64 and 3.24, whereas *csaI* was downregulated 5.74 and 6.06, respectively. Through qPCR analysis, we discovered that *csaR* and *csaI* were both downregulated at least 10-fold in PA23-6863 and PA23*phzR* (Fig. 3). In the case of *aurI* and *aurR*, differential expression could not be determined from the RNAseq data due to low read coverage of this region. Nevertheless qPCR analysis revealed that *aurI* was upregulated, while *aurR* was downregulated in PA23-6863 (Fig. 3). A similar trend was seen in PA23*phzR*, however expression of *aurI* and *aurR* was not significantly different from PA23 (Fig 3). While the regulatory network governing *aurIR* and *csaIR* expression has not yet been determined, this hierarchical arrangement is similar to that observed in *P. aeruginosa* which employs two AHL-based QS systems called Las and Rhl. In this bacterium, both the transcriptional activator (*rhlR*) and the AHL synthase (*rhlI*) genes are under control of the Las system (50, 51).

### Regulatory genes showing differential expression in PA23*phzR* or PA23-6863

Twelve genes encoding regulators belonging to diverse families including AraC, ArsR, DeoR, GntR, LysR and MarR, exhibited altered expression exclusively in PA23*phzR* (Table 4). The AHL-deficient strain displays differential expression of 18 other regulatory genes (Table 4). This latter group includes two homologues of MvaT and MvaV, which are functionally and structurally similar to the H-NS family of regulators reported to play a role in exoproduct secretion by biocontrol and pathogenic pseudomonads (12, 52). In PA23-6863, *mvaT* and *mvaV* exhibited a 1.73- and 2.92-log_2_ fold reduction in gene expression, respectively. In *P. protegens* CHA0, biocontrol activity against *Pythium ultimum* was virtually abolished in *mvaV mvaT* double mutants, and reduced in *mvaT* and *mvaV* single mutants (52). Surprisingly, MvaT and MvaV are repressors of most genes encoding exoproducts such as DAPG, HCN and exoproteases in CHA0, while positively modulating the production of PLT and siderophores. In *P. aeruginosa* PAO1, MvaT is a global regulator of virulence factors and biofilm formation, and is involved in transcriptional repression of QS (53). At present, the role of the aforementioned regulators, including MvaT and MvaV, in PA23 physiology has yet to be defined.

## Summary

Exploration of the QS regulon of biocontrol strain PA23 has revealed that approximately 13% of the genome is under QS control. This circuitry regulates diverse aspects of PA23 physiology that extend well beyond the secreted factors required for fungal antagonism. We believe that much of the QS regulon is subject to indirect control as *phz*-box elements were identified upstream of only a small percentage of QS-regulated genes. The fact that numerous transcriptional regulators show altered expression in the absence of QS further supports this notion. The number of differentially expressed genes that are unique to the AHL-deficient strain compared to the *phzR* mutant suggests that the Csa and/or Aur regulons are quite expansive and likely govern more than cell surface properties (14). Future transcriptomic analysis of *csaRI*- and *aurRI*-mutants should be conducted to reveal the scope of genes under CsaRI and AurRI QS control. Such studies will undoubtedly uncover interactions with other regulators, adding another layer to the increasingly complex cascade governing expression of PA23 biocontrol factors.

## Acknowledgements

We thank D. Khan for technical assistance and Dr. D. Haas for plasmid pME6863. This work was supported by research grants from the Natural Sciences and Engineering Research Council (NSERC) discovery grants program (T.R. de K. and M.F.B). M. Becker was supported by an NSERC Vanier Scholarship.

## Supporting information

**S1 Fig.** Autoinducer produced by PA23 and derivative strains, assessed using the AHL biosensor strain, *Chromobacterium violaceum* CVO26. Picture is representative of five biological replicates obtained from three independent experiments.

**S2 Fig.** Differentially expressed genes in (A) PA23*phzR* and (B) PA23-6863 when compared to PA23 wild type. Significantly differentially expressed genes are divided into functional categories based on Cluster of Orthologous Groups (COGs) (54). Genes important for the synthesis and regulation of biocontrol products are marked.

**S3 Fig.** Motility of PA23, PA23*phzR* and PA23-6863. A) Motility genes upregulated in PA23*phzR* and PA23-6863 compared to PA23. B) Swim plates (0.3% agar) after 24, 48, and 72 h of incubation; Swarm plates (0.8% agar) after 96 h incubation. Pictures are representative of five biological replicates obtained from three independent experiments.

**S1 Table.** RNA-sequencing library reads mapped to the *Pseudomonas chlororaphis* PA23 genome.

**S2 Table.** Differentially expressed genes in PA23*phzR* compared to PA23 wild type.

**S3 Table.** Differentially expressed genes in PA23-6863 compared to PA23 wild type.

